# ZEISS arivis Cloud: a cloud-based platform for deep learning model training and scalable bioimage analysis

**DOI:** 10.64898/2026.08.07.743540

**Authors:** Sreenivas Bhattiprolu, Manita Toor, Sebastian Soyer

## Abstract

Modern biological imaging generates large, complex datasets that require scalable and reproducible image analysis methods. Deep learning has demonstrated strong performance on bioimage segmentation tasks, but training custom models has remained inaccessible to many researchers due to requirements for GPU infrastructure, programming expertise, and large annotated training datasets. ZEISS arivis Cloud is a browser-based platform for deep learning model training that addresses these barriers through partial annotation support, AI-assisted labeling with SAM (Segment Anything Model), pretrained model initialization, and automatically configured training pipelines requiring no machine learning expertise. The platform supports two segmentation tasks: semantic segmentation using a U-Net-style architecture with an EfficientNet encoder and PixelShuffle decoder, and instance segmentation based on Mask2Former with a Swin-Tiny backbone. Both pipelines incorporate microscopy-specific adaptations including smooth tiling, multi-channel input support, dataset-specific normalization, and partial-annotation-aware loss functions protected by patents US-20240078681-A1 and US-20250111519-A1. Trained models integrate directly with ZEISS arivis Pro for pipeline-based image analysis, ZEISS arivis Hub for parallel execution across large datasets, and ZEISS ZEN for content-aware guided acquisition. We describe the platform architecture, training methodology, segmentation architectures, reproducibility and versioning mechanisms, and FAIR compliance, and illustrate the complete workflow through two intestinal organoid imaging examples. arivis Cloud is freely accessible to student users; other users access the platform via subscription at https://www.arivis.cloud/.

## 1. Introduction

Modern biological imaging has undergone a fundamental shift in scale and complexity. Advances in fluorescence microscopy, volumetric imaging, and high-content screening have made it routine to acquire three-dimensional image datasets spanning entire tissue sections, organoid cultures, or multiwell plates. Extracting quantitative information from these datasets, however, remains a significant bottleneck. Segmentation and object classification, tasks that once required manual annotation by trained image analysts, now need to scale to thousands of images while remaining reproducible across experiments, instruments, and laboratories.

Deep learning has substantially advanced the state of the art in bioimage segmentation over the past decade [1, 2]. Models trained on annotated biological images can generalize across imaging conditions and outperform classical threshold-based and machine learning approaches on complex segmentation tasks. Yet adoption among life scientists remains uneven. Training a deep learning model has historically required access to GPU computing infrastructure, familiarity with Python-based machine learning frameworks, and the ability to annotate large training datasets, prerequisites that exclude the majority of researchers who are domain experts in biology rather than computational methods.

A second, less discussed problem is the disconnection between image analysis and the broader experimental workflow. In most laboratories, image acquisition, segmentation, and quantitative analysis are performed as sequential, largely manual steps using different software tools. Pipelines built for one experiment are rarely portable to the next. Results are difficult to reproduce because the analysis parameters, model versions, and software configurations are seldom documented with the same rigor applied to the biological experiment itself. As imaging throughput increases, these gaps in reproducibility and scalability become increasingly consequential.

ZEISS arivis Cloud was developed to address these challenges within a unified, browser-based environment that requires no local compute infrastructure or programming expertise. The platform enables researchers to annotate images, train deep learning segmentation models, and run those models to segment multidimensional images without leaving a web browser. Trained models integrate directly with ZEISS arivis Pro for pipeline-based image analysis, ZEISS arivis Hub for parallel execution across large datasets, and ZEISS ZEN for feedback-driven intelligent image acquisition. Together, these components connect model training to the full experimental workflow, from acquisition through quantitative measurement.

In this manuscript, we describe the architecture, training methodology, and deployment capabilities of ZEISS arivis Cloud. We discuss the design decisions that make deep learning model training accessible to researchers without machine learning expertise, including support for partial annotations, AI-assisted labeling, and pretrained model initialization. We also describe how trained models are deployed across the ZEISS software ecosystem and illustrate the complete workflow through two application examples drawn from intestinal organoid imaging.

## 2. The ZEISS arivis Ecosystem

ZEISS arivis Cloud is one component of a broader software ecosystem designed to support the complete bioimage analysis workflow, from image acquisition through large-scale quantitative analysis. Understanding the relationship between the components helps clarify where arivis Cloud fits and why integration across them matters for reproducible research.

ZEISS arivis Cloud is a browser-based platform for image annotation and deep learning model training. It requires no local software installation or dedicated GPU hardware, and is accessible to any researcher with a compatible web browser. Student users have free access to the platform; other users access it via subscription. Models trained in arivis Cloud can be exported and deployed across the other components of the ecosystem.

ZEISS arivis Pro is a desktop application for interactive and automated image analysis. It accepts images from virtually any microscope vendor and file format, and supports datasets of virtually unlimited size. Analysis workflows in arivis Pro are configured as pipelines, in which image processing and segmentation operations are connected in sequence to produce reproducible, automated analysis from raw images through to quantitative measurements. Deep learning models trained in arivis Cloud can be imported directly into these pipelines, alongside classical segmentation methods and third-party models such as Cellpose [3].

ZEISS arivis Hub is a server-based platform for executing arivis Pro pipelines at scale. Once a pipeline has been developed and validated on a representative sample in arivis Pro, it can be deployed to arivis Hub for parallel processing across large collections of images, such as multiwell plate experiments or longitudinal datasets. Results are aggregated and accessible through a web-based interface, including heatmap visualizations for high-content screening applications.

ZEISS ZEN is the acquisition and control software used across ZEISS microscope systems. ZEN supports guided acquisition workflows in which image analysis, including deep learning segmentation, is performed during acquisition to identify regions of interest and direct subsequent imaging steps. This closes the loop between model inference and microscope control, enabling content-aware acquisition strategies without manual intervention.

Together, these four components form an integrated workflow in which a model trained once in arivis Cloud can be applied at the point of acquisition in ZEN, embedded in a reproducible analysis pipeline in arivis Pro, and executed across entire experimental datasets in arivis Hub (Figure 1).

**Figure 1.**
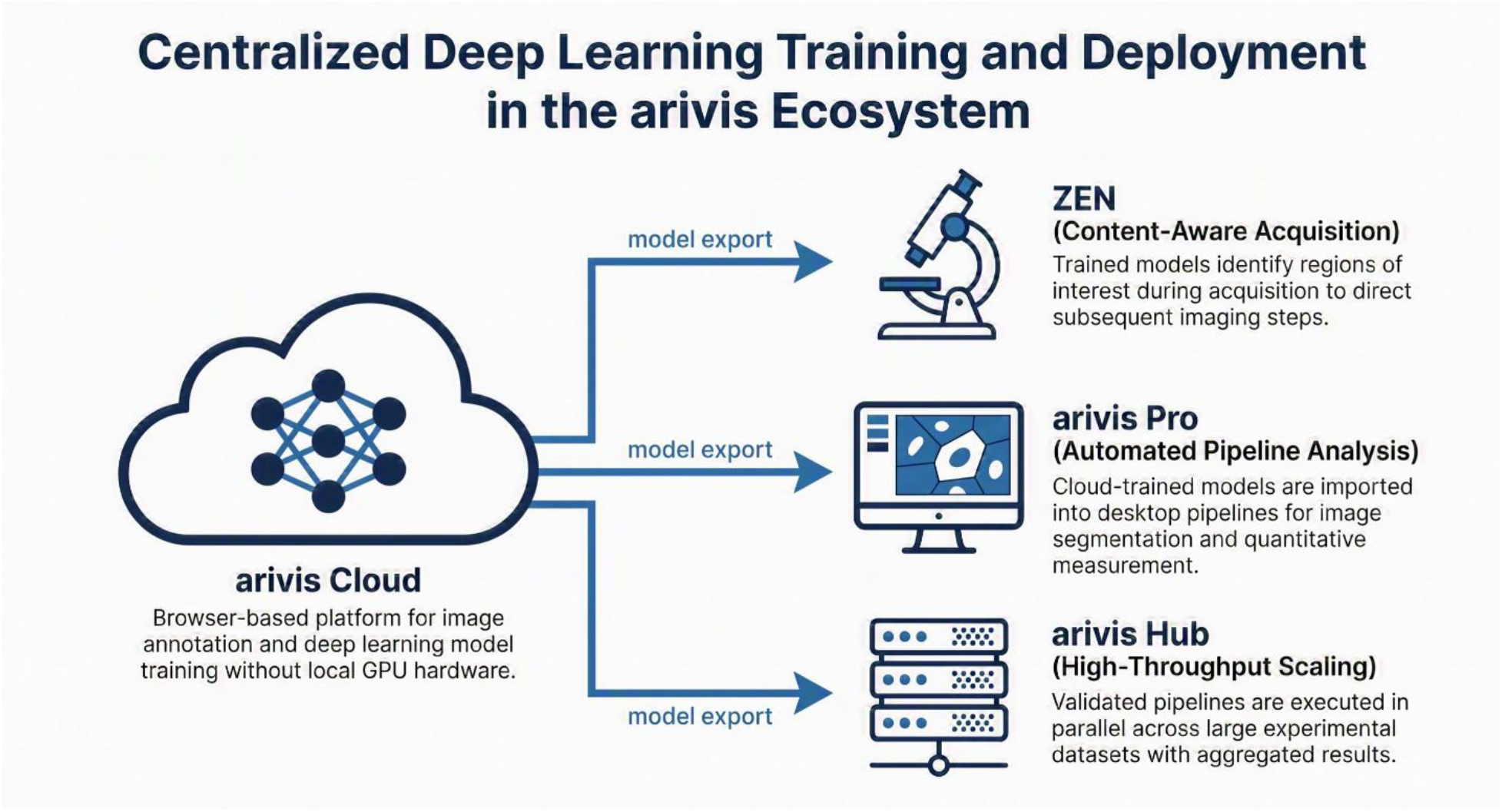
Centralized deep learning training and deployment in the arivis ecosystem. ZEISS arivis Cloud serves as the central model training hub, exporting trained models to three downstream components: ZEISS ZEN for content-aware guided acquisition, ZEISS arivis Pro for automated pipeline-based image analysis, and ZEISS arivis Hub for high-throughput parallel processing across large experimental datasets.

**Figure 2.**
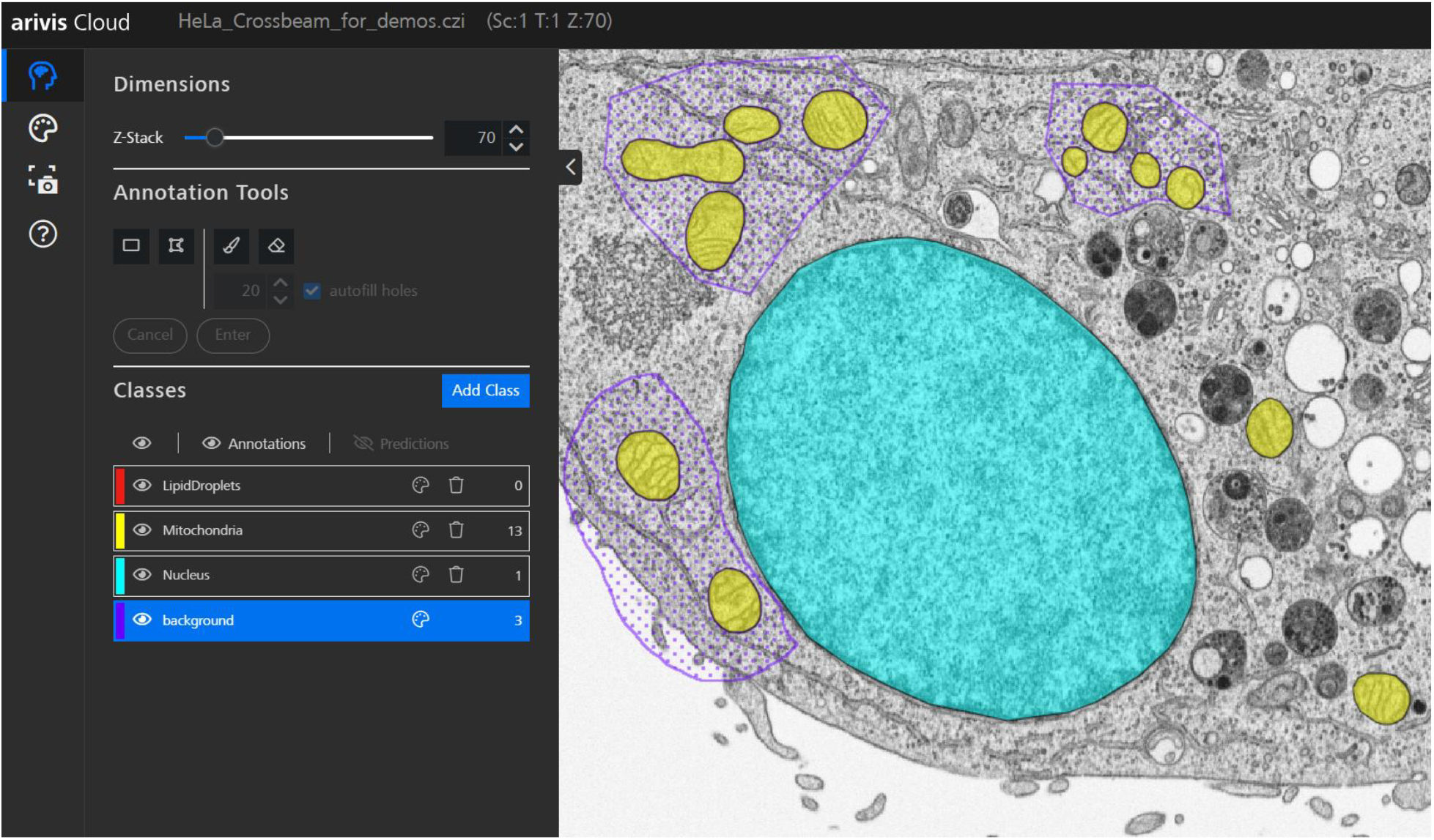
Annotation interface in ZEISS arivis Cloud showing partial annotations for mitochondria (yellow) and background (purple) classes on a FIB-SEM slice of a HeLa cell. Partial annotations allow users to label only the most informative regions rather than every pixel in the training image. Sample courtesy of Anna Steyer and Yannick Schwab, EMBL.

## 3. Platform Architecture and Access

ZEISS arivis Cloud operates as a cloud-native, browser-based application hosted on Microsoft Azure. Users access the platform through a standard web browser without installing local software or configuring GPU hardware. All compute for model training and inference is handled server-side, making the platform accessible on any device capable of running a modern browser, including laptops without dedicated graphics hardware.

The platform supports the most widely used microscopy image formats, including CZI, OME-TIFF, TIFF, PNG, and JPG, among others. Images are stored internally in the CZI format, which is the native file format used across ZEISS imaging systems. Researchers working with ZEISS instruments can therefore upload their data directly without any conversion step. Images acquired on non-ZEISS instruments or stored in other formats, including OME-TIFF, TIFF, PNG, and JPG, are converted to CZI upon import. In practice, most proprietary microscopy software supports export to open formats such as TIFF or PNG, and since model training does not require extremely large datasets, researchers can select and upload a representative subset of images rather than transferring entire acquisition archives.

Access to arivis Cloud is tiered by user type. Student users receive free access to the platform, supporting use in academic training and coursework. Non-student researchers and institutions access the platform via subscription. Authentication is managed through ZeissID, a unified identity system used across ZEISS digital products.

Although the examples presented in this manuscript are drawn primarily from biological imaging, the platform is application-agnostic and is used across a broad range of scientific and industrial domains, including materials science, geoscience, and electronics inspection.

Data and model retention follows a user-controlled policy. Uploaded datasets, annotations, and trained model artifacts are retained on the platform until the user explicitly deletes them or closes their account. This gives researchers control over the lifecycle of their training data and models, which is relevant for long-running projects and reproducibility requirements.

## 4. Annotation and Training: Reducing the Expert Bottleneck

### 4.1 The annotation challenge in biological imaging

Training a deep learning segmentation model requires labeled examples from which the model learns to distinguish structures of interest from background. In biological imaging, generating these labels is non-trivial. Microscopy images are often large, three-dimensional, and contain structures whose boundaries are ambiguous or context-dependent. Traditional supervised learning approaches require dense, pixel-wise annotation of entire training images, a process that is time-consuming even for experienced annotators and that scales poorly with image size and dataset complexity.

ZEISS arivis Cloud addresses this through several complementary mechanisms that reduce the annotation effort required to train a reliable model, while maintaining the flexibility researchers need to define biologically meaningful classes.

### 4.2 AI-assisted annotation with SAM

The platform integrates SAM (Segment Anything Model) [4] to accelerate the annotation process. SAM is a promptable segmentation model that can generate object masks from minimal user input such as clicks or bounding boxes. Within arivis Cloud, researchers can use SAM-generated suggestions as starting points for annotation, correcting or accepting them interactively. This substantially reduces the time required to label individual objects compared to manual pixel painting, particularly for instance segmentation tasks where individual object boundaries must be precisely delineated.

### 4.3 Partial annotations

A key design principle of arivis Cloud is that researchers do not need to annotate every pixel in every training image. The platform supports partial annotations, in which only a subset of regions within a training image are labeled. Unlabeled regions are not treated as negative examples during training. Instead, the loss function is modified to restrict supervision to regions that have been explicitly annotated, allowing the model to learn from incomplete labels without being penalized for correct predictions in unannotated areas.

For semantic segmentation, this is implemented through masked pixel supervision, in which a weight map identifies annotated regions and confines the gradient signal accordingly. For instance segmentation, the Mask2Former-based pipeline uses a partial-annotation-aware training setup in which loss computation is focused on reliably annotated regions. The approach underlying these implementations is protected by patents US-20240078681-A1 and US-20250111519-A1. In practice, researchers can begin training with as few as approximately 20 annotated objects and iteratively refine the model by adding annotations to regions where the current model struggles, following a data-centric rather than model-centric development strategy (Figure 3).

**Figure 3.**
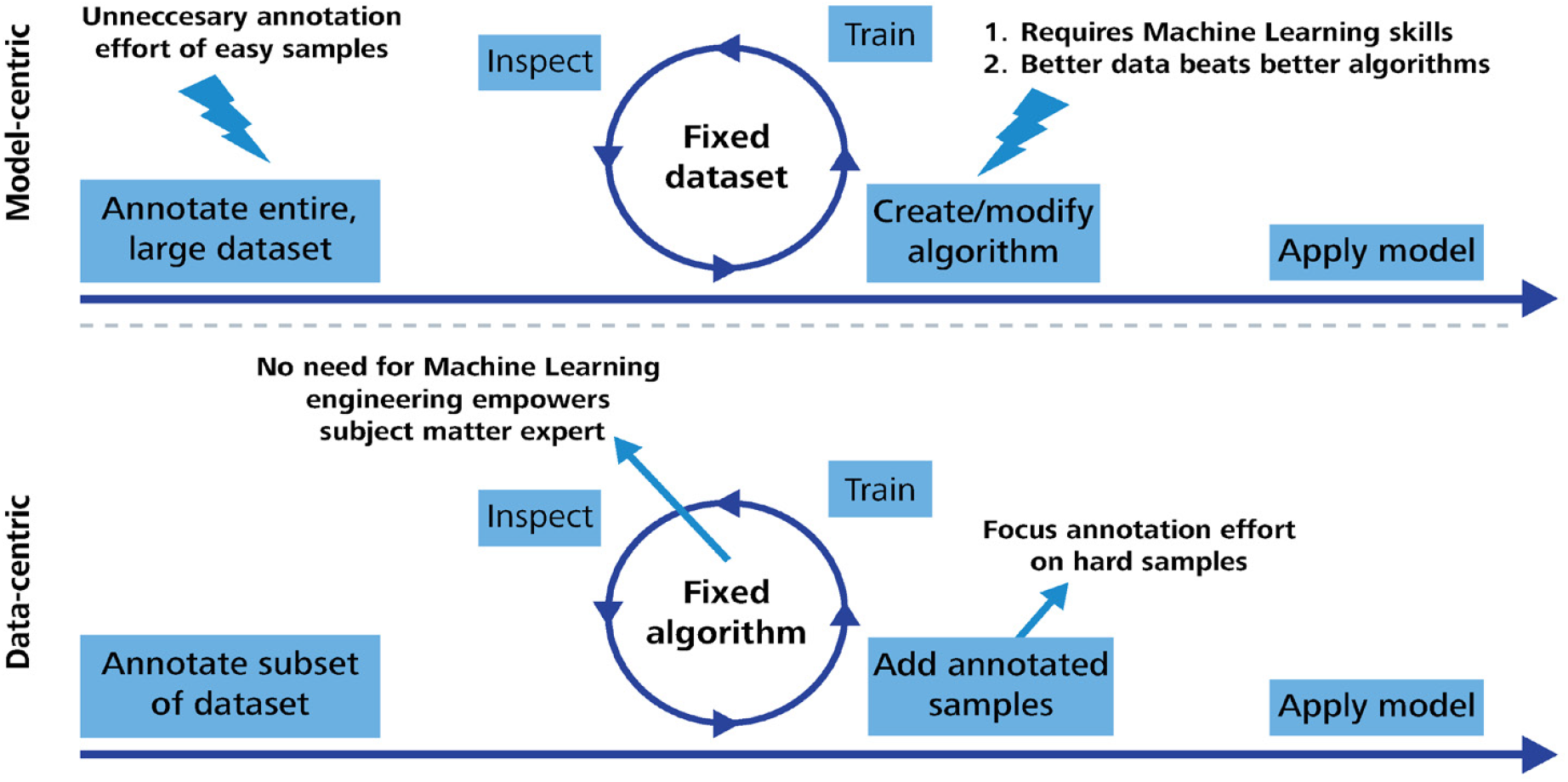
Comparison of model-centric and data-centric approaches to deep learning model development for image segmentation. In the model-centric approach (top), a large fixed dataset is annotated and the algorithm is iteratively modified to improve performance. In the data-centric approach (bottom), the algorithm is fixed and the training dataset is iteratively expanded with targeted annotations focused on challenging regions, reducing overall annotation effort.

**Figure 4.**
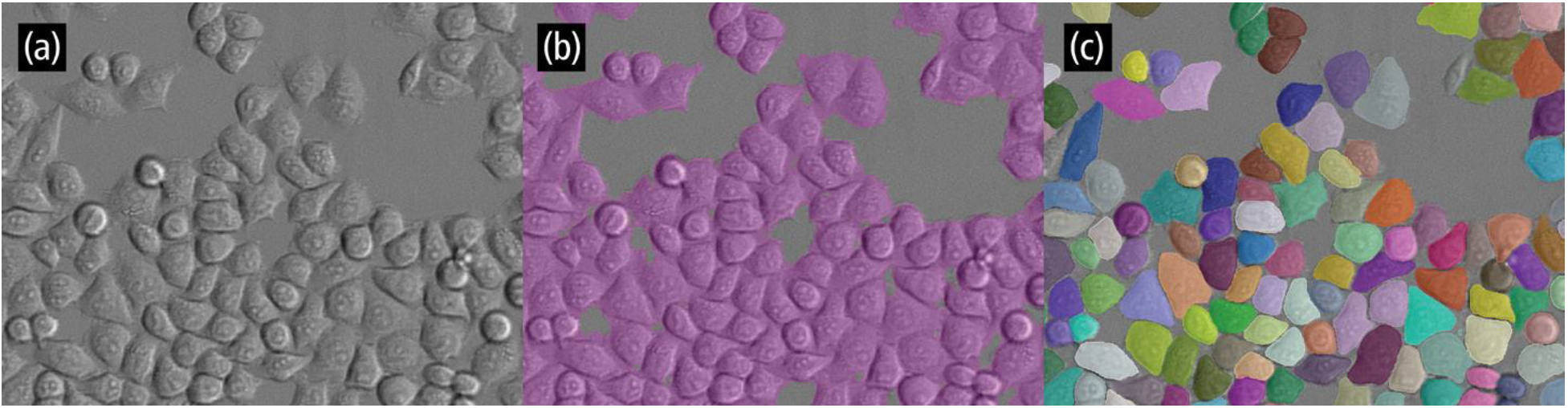
Comparison of semantic and instance segmentation approaches applied to phase contrast cell images. (a) Original phase contrast image. (b) Semantic segmentation result in which all pixels corresponding to cells are assigned to a single class (purple), without distinguishing individual cells. (c) Instance segmentation result in which each individual cell is delineated as a distinct object (shown in random colors), enabling object-level quantification even for touching cells.

### 4.4 Pretrained weights and transfer learning

Both segmentation pipelines support pretrained weight initialization to reduce the amount of labeled data needed during training. For 2D semantic segmentation, the EfficientNet encoder is initialized from ImageNet weights [5]. The instance segmentation model uses a Swin-Tiny backbone also initialized from pretrained weights [6]. It should be noted that the 3D semantic segmentation mode does not currently use pretrained initialization and trains from random weight initialization. No microscopy-specific public pretraining was used in any pipeline; the domain adaptation to microscopy data is achieved through the training process itself, supported by the mechanisms described in this section.

### 4.5 Image augmentation

To improve model generalization across imaging conditions, the training pipelines apply image augmentation automatically during training. Augmentations are selected to be appropriate for microscopy data and include geometric transforms such as flips and rotations, random crops and resizing, and intensity-based transforms including blur and noise. These augmentations are applied in the background without requiring configuration by the user, exposing the model to a broader range of image appearances than the annotated training set alone would provide.

### 4.6 Automatic boundary annotation

For instance segmentation tasks, annotating the precise boundary between adjacent objects and background is one of the most demanding aspects of labeling. arivis Cloud simplifies this by automatically defining boundary annotations from user-provided object labels, reducing the manual effort required to achieve clean boundary supervision.

### 4.7 Data-centric training strategy

The training philosophy underlying arivis Cloud follows a data-centric rather than model-centric approach (Figure 3). In a model-centric workflow, the training dataset is fixed and the algorithm is iterated to improve performance. In a data-centric workflow, the algorithm is fixed and the training data is iteratively improved by adding annotations to challenging or underrepresented regions. This is particularly well suited to microscopy applications, where imaging conditions vary substantially between experiments and where the diversity of the training dataset is often the limiting factor in model generalization. The platform provides the annotation tools needed to implement this strategy efficiently, allowing researchers to inspect current model predictions, identify failure cases, and add targeted annotations rather than labeling uniformly across the dataset.

## 5. Segmentation Architectures

ZEISS arivis Cloud supports two distinct segmentation tasks, each implemented with a dedicated deep learning architecture chosen and adapted for microscopy image data.

### 5.1 Semantic segmentation

Semantic segmentation assigns each pixel in an image to one of a set of user-defined classes. This is appropriate for tasks such as identifying tissue regions, classifying cellular compartments, or separating foreground structures from background where individual object identity is not required.

The semantic segmentation pipeline uses a U-Net-style encoder-decoder architecture [2] with an EfficientNet encoder [5] and a PixelShuffle-based decoder [7]. EfficientNet provides an efficient and scalable feature extraction backbone that balances parameter count with representation capacity. PixelShuffle upsampling in the decoder avoids the checkerboard artifacts associated with transposed convolution upsampling, producing cleaner segmentation boundaries. The architecture supports multi-channel inputs, allowing images with any number of fluorescence channels to be used directly without modification. Both 2D and 3D operation modes are supported. Normalization is computed from dataset statistics rather than fixed values, adapting the model to the intensity distributions of the specific imaging data being analyzed.

Default tile sizes for semantic segmentation are 1024 x 1024 pixels, selected to provide sufficient spatial context for most microscopy segmentation tasks. Tile size adapts based on image characteristics where necessary. One practical constraint is that the tile size cannot exceed the smallest side of the smallest image in the training set; users working with small images should be aware that this may result in a reduced effective tile size.

### 5.2 Instance segmentation

Instance segmentation identifies and delineates individual object instances within an image, assigning a unique label to each detected object. This is required for tasks where object-level measurements are needed, such as counting cells, measuring individual organoid volumes, or extracting morphological parameters from each nucleus independently.

The instance segmentation pipeline is based on Mask2Former [8], a transformer-based architecture that formulates instance segmentation as a set prediction problem. Mask2Former uses a Swin-Tiny backbone [6] for feature extraction. Microscopy-specific adaptations include flexible multi-channel input handling, dataset-specific normalization, and a partial-annotation-aware training loss as described in section 4.3. Following model inference, a non-maximum suppression step removes duplicate detections. Tile merging for instance segmentation is described in section 5.3. Default tile sizes for instance segmentation are 320 x 320 pixels.

### 5.3 Tiling strategies for large images

Both pipelines process large images by dividing them into overlapping tiles, running inference on each tile independently, and merging the results. The strategies used to combine tile predictions differ between the two pipelines.

For semantic segmentation, arivis Cloud uses a smooth tiling approach in which predictions from overlapping tiles are blended by weighting pixels according to their distance from the tile edge. Pixels near the center of a tile receive higher weight than pixels near the edge, reflecting the greater spatial context available to the model at the tile center. This produces seamless segmentation results across large images without visible boundary artifacts (Figure 5).

**Figure 5.**
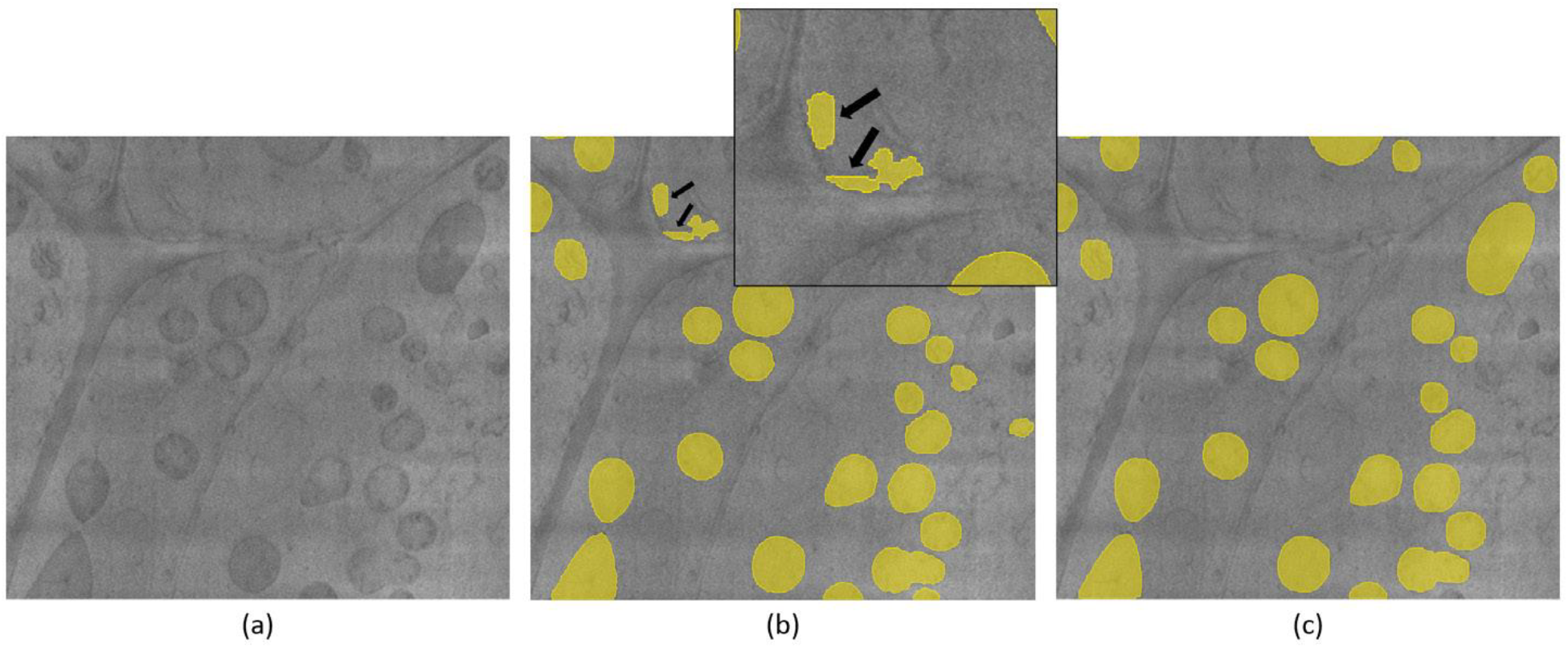
(a) Cryo-electron microscopy image of a cell showing mitochondria. A Deep Learning model has been trained using ZEISS arivis Cloud to segment faint mitochondria from the background. (b) The segmentation result without smooth blending produces noticeable artifacts along the patch edges, leading to incorrect classification of edge pixels as mitochondria. Improbably straight edges are also clearly visible, as indicated by the black arrows. (c) The seamless integration of patches using smooth tiling creates a segmented image without any visible anomalies. Sample courtesy of Dr. York-Dieter Stierhof from Eberhard Karl University of Tübingen.

For instance segmentation, tile merging operates at the object level rather than the pixel level. Following inference on individual tiles, a proprietary algorithm reconciles object instances detected in neighboring tiles, resolving overlapping or split detections across tile boundaries to produce a consistent set of object instances across the full image.

### 5.4 Fixed hyperparameters and microscopy-specific tuning

A deliberate design decision in arivis Cloud is that model hyperparameters are not exposed to the user for manual tuning. Learning rates, augmentation schedules, batch sizes, and architectural settings have been configured and validated for microscopy image data by the development team. This reflects the intended user base: domain experts across a wide range of scientific and industrial fields who should not be required to understand the implications of hyperparameter choices to obtain a reliable segmentation model. arivis Cloud is application-agnostic and is used across biological imaging, materials science, geoscience, electronics inspection, and other fields where image segmentation plays a role. The training methodology and architecture decisions described in this manuscript apply equally across these domains. Additional training considerations include class and border weighting and postprocessing tailored to object-style outputs, all applied automatically as part of the training pipeline.

## 6. Model Evaluation and Reproducibility

### 6.1 Training metrics

During training, arivis Cloud tracks metrics to characterize model performance. For semantic segmentation, the primary validation metric is pixel-wise intersection over union (IoU), reported both as a mean across all classes and on a per-class basis. Loss curves are tracked throughout training and training artifacts include a history file recording metric progression across epochs.

The platform is designed for researchers who are domain experts rather than machine learning practitioners. Accordingly, raw training metrics are not prominently surfaced in the user interface, as interpreting loss curves and IoU values requires familiarity with deep learning training dynamics that many users do not have. Researchers who require access to training metrics for publication purposes can retrieve the stored training artifacts, which include JSON configuration files, metric logs, and model checkpoint files.

The train/validation split strategy differs between the two pipelines. For semantic segmentation, an automatic 80:20 split is applied at the tile level during training. For instance segmentation, the full annotated dataset is used for training without a held-out validation split, as the small annotation sets typical in practice make withholding data impractical.

### 6.2 Reproducibility and versioning

Reproducibility in deep learning-based image analysis depends on the ability to recover not just a trained model but the complete set of conditions under which it was produced. arivis Cloud addresses this through systematic versioning of training inputs and outputs.

Each training run is assigned a unique run identifier. Dataset identifiers are associated with each run, providing a traceable link between a trained model and the images and annotations used to produce it. Training hyperparameters, including learning rate, augmentation configuration, tile and patch sizes, batch size, and model architecture settings, are stored in JSON-format configuration files and logged with each run. Trained model artifacts, including weight files and model configuration, are stored with model name and version identifiers that allow a specific model version to be unambiguously referenced.

Within the platform, trained models can be shared with collaborators and applied to new data using the stored model artifact and configuration, ensuring that inference conditions are identical to those used during development. This combination of dataset versioning, hyperparameter logging, and model versioning provides the documentation needed to describe an analysis pipeline in a methods section and for others to understand the conditions under which results were produced.

## 7. FAIR Compliance

The FAIR principles, requiring that research outputs be Findable, Accessible, Interoperable, and Reusable [9], have become an important framework for evaluating research software and data management practices in the life sciences. We describe here how arivis Cloud addresses each principle.

### Findable

The platform assigns unique identifiers to datasets, trained models, model versions, and training runs. These identifiers serve as persistent internal references that allow specific training configurations and model versions to be located and retrieved. A model registry within the platform allows users to search their trained models by name.

### Accessible

The platform is accessible through any standard web browser following authentication via ZeissID. Student users have free access; other users access the platform via subscription. The authentication protocol is well-defined and consistently enforced. Data and models are retained until explicitly deleted by the user, ensuring that resources remain accessible for the duration of a research project.

### Interoperable

arivis Cloud accepts images in the most widely used microscopy formats, including CZI, OME-TIFF, TIFF, PNG, and JPG, among others, covering data from both ZEISS and non-ZEISS acquisition systems. Trained semantic segmentation models are exported in the CZANN/CZMODEL format for use in downstream ZEISS software. Trained instance segmentation models are packaged as runnable containers retrievable via the ZEISS AI registry and deployable within the ZEISS software ecosystem. Segmentation results produced within the platform can be retrieved as label masks. Training metadata, including dataset properties, class definitions, normalization statistics, and hyperparameter configurations, are stored as machine-readable JSON files alongside trained model artifacts.

### Reusable

Trained models can be shared within the platform by model identifier, allowing collaborators to apply an identical model to new data. The complete training configuration stored with each model provides the metadata needed to understand the conditions under which the model was produced and to assess its suitability for reuse on new datasets. The platform does not currently provide a community model repository, but the model sharing capability within institutional or collaborative accounts supports reuse within research groups and consortia.

## 8. Deployment and Workflow Integration

A model trained in arivis Cloud is most useful when it can be applied within the broader context of an experimental workflow. The ZEISS arivis ecosystem provides three distinct deployment pathways, each addressing a different stage of the imaging and analysis pipeline.

### 8.1 Pipeline-based analysis in arivis Pro

ZEISS arivis Pro is a desktop image analysis application that accepts images from virtually any microscope vendor and file format and supports datasets of virtually unlimited size. Image analysis in arivis Pro is organized around the pipeline concept, in which individual processing and segmentation operations are connected in sequence. Data flows automatically from one step to the next, and the complete pipeline can be saved, shared, and re-applied to new datasets to ensure reproducible analysis.

Trained models from arivis Cloud can be imported directly into arivis Pro pipelines as segmentation operations. They can be combined with classical segmentation methods, morphological operations, object filtering, and measurement extraction within the same pipeline. Third-party models, including Cellpose [3], are also supported within this framework, allowing researchers to use the most appropriate segmentation approach for each structure of interest within a single automated workflow. Once segmentation is complete, arivis Pro extracts quantitative measurements from segmented objects, including morphological features such as volume, surface area, sphericity, compactness, and equivalent ellipse parameters, as well as per-channel intensity statistics, texture features, and relational measurements such as inter-object distances. Custom features can also be defined. The pipeline from raw image to measurement table is fully automated and documented by the saved pipeline configuration.

### 8.2 Scaled execution in arivis Hub

Once a pipeline has been developed and validated in arivis Pro on a representative subset of images, it can be deployed to ZEISS arivis Hub for execution across large image collections. arivis Hub is a server-based platform that runs arivis Pro pipelines in parallel across multiple datasets, with the degree of parallelism determined by the number of available compute workers. This makes it practical to apply a validated analysis pipeline to entire multiwell plate experiments, longitudinal datasets, or large cohort studies without rewriting or reconfiguring the pipeline.

Results from Hub runs are aggregated and accessible through a web-based interface. For high-content screening applications, arivis Hub provides heatmap visualizations that display per-well summary statistics across a plate layout, enabling rapid identification of experimental conditions that differ from controls. Per-object and per-image data tables are available for download for downstream statistical analysis.

### 8.3 Intelligent acquisition in ZEN

ZEISS ZEN is the acquisition and control software used across ZEISS imaging systems. ZEN supports guided acquisition workflows that integrate image analysis directly into the acquisition process. In a guided acquisition workflow, an initial low-magnification overview scan is followed by automated image analysis to identify objects or regions of interest, and the microscope then acquires high-resolution, multi-dimensional images of only those identified positions. This approach makes efficient use of microscope time and ensures that high-resolution imaging resources are directed toward biologically relevant targets.

Deep learning models trained in arivis Cloud can be deployed within ZEN guided acquisition workflows to perform the object detection step. This is particularly valuable when the structures of interest are difficult to identify by simple intensity thresholding due to morphological complexity, clustering, or variable positioning within the field of view. The integration of arivis Cloud models into ZEN closes the loop between model training and intelligent acquisition, enabling content-aware imaging strategies that adapt to the biological content of each sample.

## 9. Application Examples

We present two application examples that illustrate the complete workflow from model training in arivis Cloud through deployment in the broader ecosystem. Both examples use intestinal organoids as the biological system, reflecting the growing importance of three-dimensional organoid culture models in developmental biology and drug discovery research.

### 9.1 Content-aware guided acquisition in ZEN

Multiwell plate experiments involving three-dimensional organoids present a practical challenge for high-resolution imaging. Organoids vary in position, depth, and morphology across wells, and acquiring full high-resolution three-dimensional stacks of every position in a plate is time-prohibitive. Automated guided acquisition in ZEN addresses this by directing high-resolution imaging only to positions containing objects that meet user-defined criteria.

In this example, murine Lgr5+ intestinal organoids were cultured in a 24-well plate and imaged on a ZEISS Celldiscoverer 7. The guided acquisition workflow consisted of three stages (Figure 6). First, a low-magnification widefield overview scan was performed across the plate to capture the distribution of organoids within each well. Second, image analysis identified individual organoids as regions of interest. For well-separated organoids with sufficient contrast, this detection step can be accomplished using intensity thresholding. In cases where organoids are clustered together or positioned near well edges, a deep learning model trained in arivis Cloud provides more robust detection by learning to discriminate organoids from background based on morphological features rather than intensity alone. Third, the microscope automatically acquired high-resolution, multi-channel three-dimensional stacks at each identified organoid position using the Airyscan detector at 20x magnification, capturing nuclei and E-cadherin channels across the full organoid depth.

**Figure 6.**
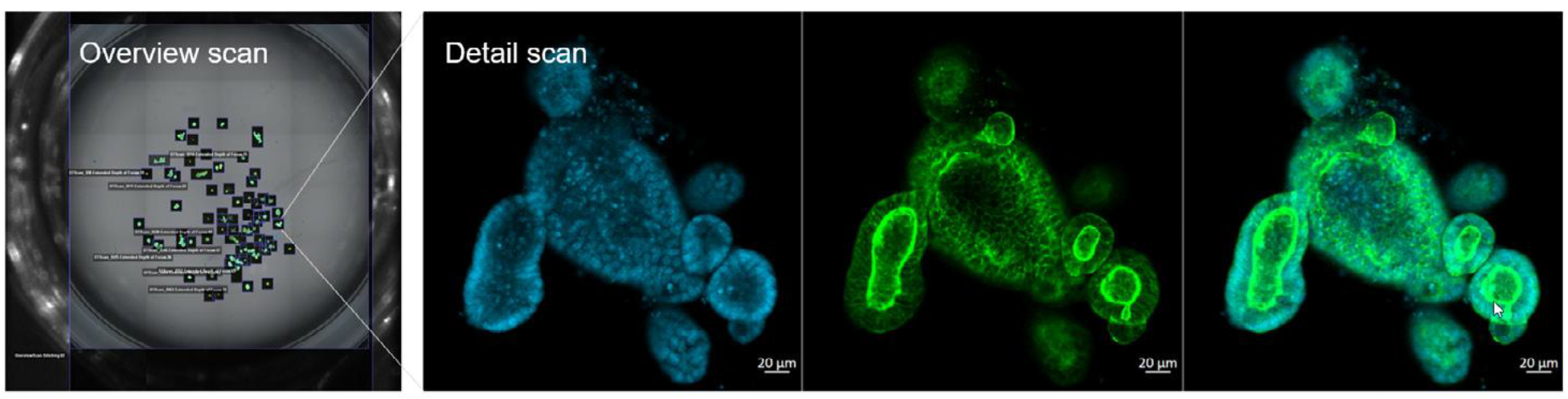
Guided acquisition of mouse Lgr5+ gut organoids in ZEN. (a) The initial low-magnification overview scan of a single well from a 24-well plate, revealing multiple organoids. The three images on the right display high-resolution two-dimensional (2D) sections of a select organoid from panel (a), highlighting nuclei (blue) in panel (b), E-cadherin (green) in panel (c), and both channels combined in panel (d). The multi-dimensional characterization of organoids within a complex 3D system is showcased. The organoids were mounted in a 3D matrix (Matrigel) and imaged on a Celldiscoverer 7. Sample courtesy of Dr. M. Lutolf, EPFL, Switzerland.

This workflow substantially reduces the imaging time required for a multiwell organoid experiment while ensuring that high-resolution data are collected for a statistically representative sample of organoids across each well. Sample courtesy of Dr. M. Lutolf, EPFL, Switzerland; imaging by Frank Vogler, ZEISS customer center.

### 9.2 Scalable organoid analysis from model training to plate-level results

The second example demonstrates the complete analysis workflow from model training in arivis Cloud through pipeline execution in arivis Pro and scaled processing in arivis Hub, using intestinal organoids to study the role of Wnt signaling in organoid formation [10].

Intestinal organoids consist of a single layer of epithelial cells surrounding a hollow lumen. Quantitative analysis of organoid morphology and cellular composition requires accurate segmentation of three distinct structures: the organoid lumen, individual nuclei, and cell bodies. These structures differ substantially in their imaging characteristics, and no single segmentation approach is optimal for all three.

For nucleus segmentation, the Cellpose algorithm [3] was used directly without custom training, as it provides reliable nucleus detection in fluorescence images across a broad range of cell types and imaging conditions. For lumen segmentation, a custom model was trained in arivis Cloud. The lumen presents a more demanding segmentation challenge due to its irregular morphology and variable contrast, and the machine learning approach enabled by arivis Cloud produced superior results compared to intensity thresholding, particularly for organoids in which lumen boundaries are indistinct. An analysis pipeline was then constructed in arivis Pro combining the Cellpose nucleus segmentation and the arivis Cloud lumen model with morphological operations, object filtering, and measurement extraction. The pipeline extracted morphological measurements including organoid volume, cell layer volume, lumen volume, and organoid roundness, as well as per-nucleus and per-cell intensity measurements for differentiation markers. Cell bodies were modeled by region growing from nucleus objects, and nuclei were stratified into cell layer and luminal compartments based on spatial relationships to the lumen segmentation.

The pipeline was first developed and validated on a representative subset of 30 organoids per experimental condition. Once validated, it was deployed to arivis Hub for parallel execution across 60 organoid datasets in a single batch run (Figure 7). Aggregated results were visualized as a plate-level heatmap of nucleus count per well, with per-organoid measurement tables available for download. This workflow identified significant differences in cell number and Aldolase B expression between control and Wnt-inhibited organoids, consistent with a role for Wnt signaling in organoid maturation.

**Figure 7.**
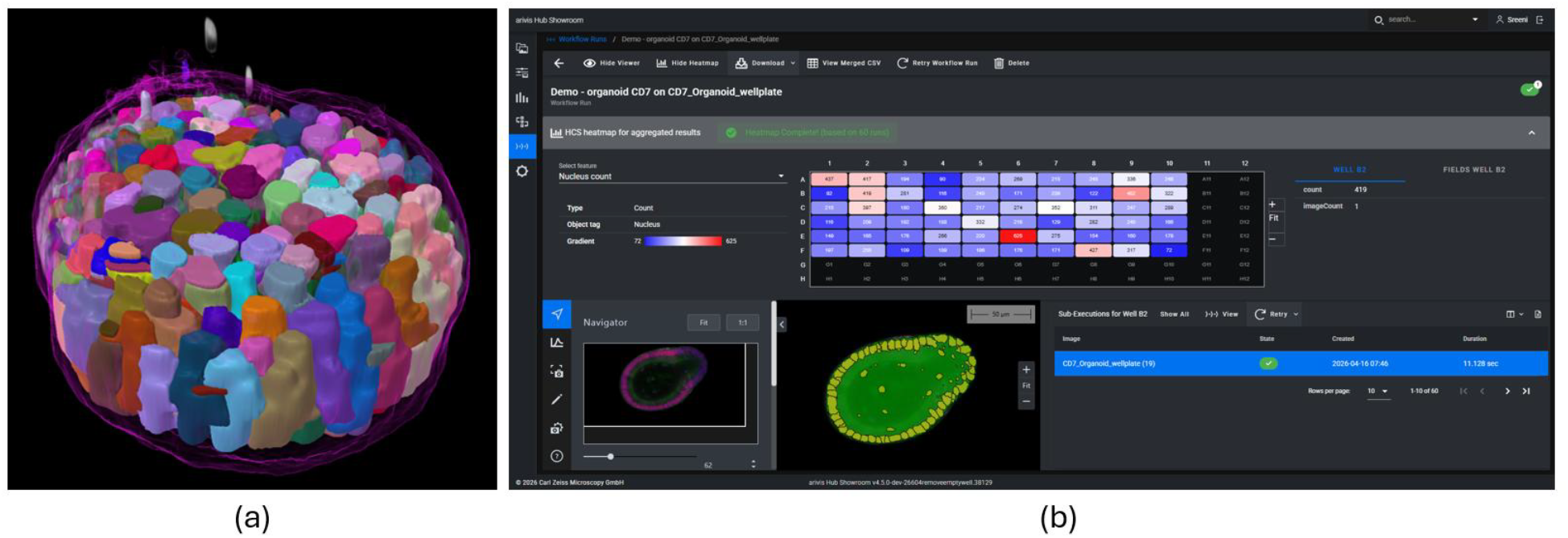
Scalable organoid analysis workflow using ZEISS arivis Cloud, arivis Pro, and arivis Hub. (a) Three-dimensional instance segmentation rendering of a murine intestinal organoid imaged at 20x magnification on a ZEISS Celldiscoverer 7, with individual cells segmented using a combination of Cellpose (nuclei) and a custom model trained in arivis Cloud (lumen). (b) arivis Hub workflow run showing aggregated nucleus count results across 60 wells of a multiwell plate, displayed as a high-content screening heatmap. The pipeline was developed and validated in arivis Pro before parallel execution across all wells via arivis Hub. Sample courtesy of Lutolf lab; imaging by Frank Vogler, ZEISS customer center.

The key design principle illustrated by this example is the separation between model development and scaled execution. The analysis pipeline was defined and validated once, then applied identically across all datasets through arivis Hub, ensuring that results are not affected by analyst-to-analyst variation or incremental changes in processing parameters across the dataset.

## 10. Discussion

ZEISS arivis Cloud addresses a practical gap in the bioimage analysis landscape: the need for deep learning model training tools that are accessible to domain experts without machine learning expertise, and that connect directly to the downstream analysis and acquisition workflows in which those models will be used.

Several existing tools address parts of this problem. ilastik [11] provides accessible machine learning-based segmentation through a graphical interface and has been widely adopted in the biological imaging community. QuPath [12] offers deep learning integration within a digital pathology-focused environment. Cellpose [3] provides a generalist cell segmentation algorithm that performs well across many imaging modalities without retraining. ZeroCostDL4Mic [13] makes deep learning model training accessible through Google Colab notebooks, removing the local compute requirement. Each of these tools has established use cases and a strong user community.

arivis Cloud occupies a distinct position in this landscape by combining browser-based model training with direct integration into acquisition and scaled analysis workflows. The ability to deploy a trained model in ZEN for guided acquisition, in arivis Pro for pipeline-based analysis, and in arivis Hub for plate-scale execution within the same ecosystem is a practical advantage for researchers running high-content experiments. The partial annotation support and data-centric training philosophy reduce the labeling burden compared to approaches that require fully annotated training datasets.

There are limitations worth acknowledging. Trained models and analysis pipelines are primarily designed for use within the ZEISS software ecosystem. While semantic segmentation models are exported in the CZANN/CZMODEL format and instance segmentation models are packaged as containers, these are not currently designed for deployment in arbitrary external environments. The platform does not currently provide a community model repository, which limits the ability to share trained models publicly in a manner that would support full reproducibility for external readers of a published study. Researchers who require quantitative model performance metrics for publication can retrieve the stored training artifacts, which include metric logs and configuration files, though these are not surfaced prominently in the standard user interface given the accessibility focus of the platform.

Future development directions include expanding the range of supported segmentation tasks, improving the exposure of training metrics for users who require them for publication, and extending model interoperability beyond the ZEISS ecosystem.

## 11. Conclusion

ZEISS arivis Cloud provides a browser-based environment for deep learning model training that is designed for domain experts rather than machine learning specialists. By combining partial annotation support, AI-assisted labeling, pretrained model initialization, and automatically configured training pipelines, the platform lowers the practical barrier to training reliable segmentation models from small annotated datasets. Integration with ZEISS arivis Pro, arivis Hub, and ZEN connects model training to the full experimental workflow, enabling reproducible analysis pipelines that scale from single images to entire plate-based experiments and that can inform acquisition decisions in real time.

The platform is accessible to student users at no cost, supporting adoption in academic research and teaching. For researchers who need to cite a specific tool in their methods section, the platform provides unique identifiers for trained models and training runs, and the training configuration and hyperparameters are stored alongside model artifacts to support methods reporting. We hope this manuscript provides the technical detail needed for researchers using arivis Cloud to describe their analysis workflow accurately and completely in published work.

## References

[1] LeCun Y, Bottou L, Bengio Y, Haffner P. Gradient-based learning applied to document recognition. Proceedings of the IEEE. 1998;86(11):2278–2324.

[2] Ronneberger O, Fischer P, Brox T. U-Net: Convolutional networks for biomedical image segmentation. In: MICCAI. 2015:234–241.

[3] Stringer C, Wang T, Michaelos M, Pachitariu M. Cellpose: a generalist algorithm for cellular segmentation. Nature Methods. 2021;18(1):100–106.

[4] Kirillov A, et al. Segment Anything. In: ICCV. 2023:4015–4026.

[5] Tan M, Le QV. EfficientNet: Rethinking model scaling for convolutional neural networks. In: ICML. 2019.

[6] Liu Z, et al. Swin Transformer: Hierarchical vision transformer using shifted windows. In: ICCV. 2021:10012–10022.

[7] Shi W, et al. Real-time single image and video super-resolution using an efficient sub-pixel convolutional neural network. In: CVPR. 2016:1874–1883.

[8] Cheng B, Misra I, Schwing AG, Kirillov A, Girdhar R. Masked-attention mask transformer for universal image segmentation. In: CVPR. 2022:1290–1299.

[9] Wilkinson MD, et al. The FAIR Guiding Principles for scientific data management and stewardship. Scientific Data. 2016;3:160018.

[10] Seidel P. From image to results: organoid analysis. ZEISS Microscopy Application Note. 2021. https://www.zeiss.com/microscopy/us/applications/biotech--pharma/organoid-analysis.html

[11] Berg S, et al. ilastik: interactive machine learning for (bio)image analysis. Nature Methods. 2019;16(12):1226–1232.

[12] Bankhead P, et al. QuPath: Open source software for digital pathology image analysis. Scientific Reports. 2017;7(1):16878.

[13] von Chamier L, et al. Democratising deep learning for microscopy with ZeroCostDL4Mic. Nature Communications. 2021;12(1):2276.

